# Giant genome of the vampire squid reveals the derived state of modern octopod karyotypes

**DOI:** 10.1101/2025.05.16.652989

**Authors:** Masa-aki Yoshida, Emese Tóth, Koto Kon-Nanjo, Tetsuo Kon, Kazuki Hirota, Atsushi Toyoda, Hidehiro Toh, Hideyuki Miyazawa, Makoto Terauchi, Hideki Noguchi, Davin H. E. Setiamarga, Oleg Simakov

## Abstract

Why some animal groups retain ancestral chromosomal complements while others change significantly is a fundamental question in evolutionary genomics. Few systems exist where accumulations of chromosomal changes can be studied in the context of morphological innovation. In coleoid cephalopods (octopus, squid, cuttlefish), an ancient coleoid chromosomal rearrangement event (ACCRE) has led to a substantial increase in the chromosome number and a new set of chromosomal homologies. Compared to typical molluscan or bilaterian genomes, ACCRE has enabled the origin of many novel regulatory regions in coleoid cephalopods. However, the discrepancies between extant octopodiform (octopus, ∼30 chromosomes) and decapodiform (squid and cuttlefish, ∼46 chromosomes) karyotypes and the direction of these evolutionary changes remain unexplained. Here we provide a draft genome assembly of the vampire squid *Vampyroteuthis* sp., the largest cephalopod genome sequenced to-date (over 10 gigabasepairs). Through syntenic comparisons, we infer that this basally branching octopodiform species shows partial retention of the chromosomal complement of Decapodiformes, indicating its more ancestral state and the derived nature of the octopod karyotype. Together with the analysis of a new chromosome-level assembly of the pelagic octopod *Argonauta hians*, we identified irreversible chromosomal fusion-with-mixing events followed by inter-chromosomal translocations in octopods. We show that this secondary reduction and mixing within octopod chromosomes has enabled the origin of a more entangled genomic configuration, shedding light onto the early evolutionary transitions within the clade. Our results offer broader insights into general patterns of chromosomal evolution following large-scale rearrangement events in animal genomes.

**Significance statement:** How changes to the ancient animal synteny result in novel chromosomal homologies is difficult to dissect due to the lack of intermediate states. Here we report that the genome of *Vampyroteuthis*, one of the largest animal genomes sequenced to-date (over 11 gigabasepairs), despite its phylogenetic position within the octopodiform cephalopods, partially retains squid and cuttlefish chromosomal complement, reflecting an ancestral karyotypic state that existed at the time of divergence between these cephalopod lineages. Our findings reveal karyotype reductions through chromosomal fusions were followed by inter-chromosomal translocations in octopods, leading to a more specialized genomic and gene regulatory architecture. These data show how chromosomal fusions can act as drivers of further inter-chromosomal rearrangements in animal genomes.

## Introduction

Cephalopods (octopuses and squids) are known for their complex behaviors^1^, including problem solving, learning by observation, executing different tasks, and exceptional camouflaging ability^1^. Although similar in size to those of mammals, coleoid nervous systems are structurally distinct and have a long evolutionary history associated with advanced cognition and neural complexity.

Coleoids diverged from nautiloids in the Late Ordovician (ca. 450 Ma)^2^ (**Figure 1a**). While extant nautiloids retained their external shells, coleoids internalized or lost theirs, enabling greater maneuverability and adaptation to diverse marine environments^3,4^. Shell loss may also have driven the evolution of sophisticated neural processing leading to adaptive and complex behaviors such as rapid decision-making and camouflaging^5^. By the Late Permian to the Middle Triassic (∼260–235 Ma), coleoids had further diversified into Octopodiformes (vampire squids and octopuses) and Decapodiformes (squids and cuttlefishes)^2,6–8^ (**Figure 1a**).

**Figure 1.**
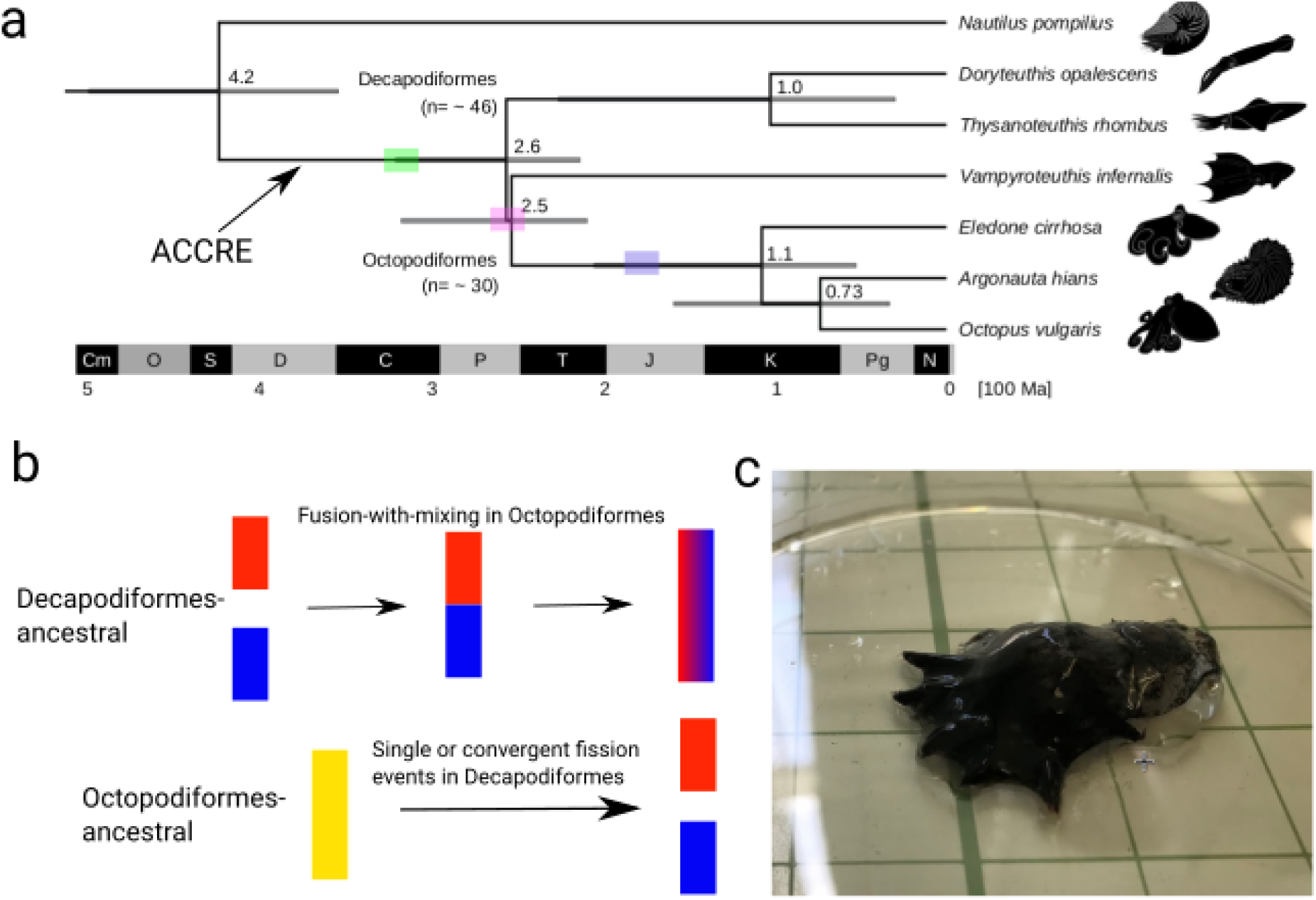
Early origins of coleoid cephalopods and conflicting chromosome evolution scenarios. (a) Phylogenetic time tree of mitochondrial genomes with the major coleoid cephalopod clades and their closest outgroups. Node branches are labelled according to the major cephalopod evolutionary transitions. (b) Two conflicting scenarios how coleoid karyotypes evolved, reduction and expansion (via fission) scenarios. Fusion-with-mixing (FWM) provides a powerful synapomorphic character to profile irreversible changes. (c) *Vampyroteuthis* specimen used for sequencing. ACCRE: ancient coleoid chromosomal rearrangement event.

The evolution of behavioral and neural complexities in coleoids, which was probably driven by clade-specific predation pressures and ecological adaptations^9–12^, likely required major genetic innovations. Recent studies have begun to uncover how large-scale changes in genome structure contributed to the evolution of their neural, cognitive, and other traits unique to coleoids. Comparative analyses show coleoids differ substantially in genome organization from both their primitive relatives and other mollusks^13,14^. While extant nautiluses retain the typical molluscan genomic structure^15–17^, coleoids underwent “ancient coleoid chromosomal rearrangement events^15^ (ACCREs)”, causing topological reorganizations that restructured gene linkages and formed novel regulatory landscapes^18^.

While ACCRE marked a key genomic transition in early coleoids, substantial lineage-specific changes between major coleoid lineages likely happened post-ACCRE. Octopuses and squids differ in chromosome numbers^15^, repeat content^19^, and expansions of gene families such as *Protocadherins*^20^ and *GPCRs*^21^. Interestingly, while coleoid chromosomes have remained largely intact within both decapodiform and octopodiform lineages, their homologies are not strictly one-to-one, with some chromosomes showing many-to-many correspondences^15^. The discrepancy between predicted typical chromosome numbers (∼30 in Octopodiformes vs. ∼46 in Decapodiformes) also raises questions about whether chromosome numbers contracted or expanded during early coleoid evolution (**Figure 1b**). However, the lack of genomic data from early branching Octopodiformes or Decapodiformes has obscured the directionality of these transitions.

The vampire squid (*Vampyroteuthis* cf. *infernalis*), a basal Octopodiformes, provides critical insight into the deep evolutionary history of coleoids^23^. This species is a globally distributed deep-sea pelagic cephalopod with an opportunistic detritivorous and zooplanktivorous feeding strategy^24,25^. This “living fossil” is the only extant representative of the vampyropods, an ancient clade within Octopodiformes (**Figure 1**) that has existed since the Mesozoic Era, from at least the Middle Jurassic (e.g., *Vampyronassa*; ca. 165 Ma) ^23,24^ with some evidence suggesting its origins in the Early Jurassic (e.g., *Teudopsis* and *Simoniteuthis*; ca.183 Ma)^26–28^. Ancient vampyropods coexisted with the ammonites and the belemnites throughout the Jurassic (∼201–145 Ma) and Cretaceous (∼145–66 Ma), occupying various ecological niches across global marine environments^27,29,30^. However, most vampyropod lineages went extinct at the end of the Cretaceous (∼66 Ma), eventually leaving extant *Vampyroteuthis* as the sole surviving species^31^. *Vampyroteuthis*’ unique phylogenetic position makes it a valuable model for resolving early genomic transition events and subsequent evolutionary trajectories in Octopodiformes and coleoids.

To enhance resolution within Octopodiformes, we also present the chromosome-level genome assembly of the muddy argonaut *Argonauta hians*, a member of Argonautoidea, an octopodiform clade composed mainly of pelagic octopods characterized by the presence of the secondarily acquired shell-like calcified eggcases^32,33^. The inclusion of these two new genomes in our analyses enables us to reveal the directionality and trajectory of chromosomal rearrangements, genomic reorganizations, and size variation across early and modern cephalopods.

## Results

We sequenced the draft genome of *Vampyroteuthis* sp. from an individual collected in the West Pacific Ocean. Genome sequencing was performed using the PacBio HiFi platform at approximately 40X coverage and assembled with hifiasm, resulting in a 14 Gb assembly. Different assembly settings were tested, yielding similar assembly sizes (**Supplementary Note**). A more stringent setting to both haplotype overestimation and to remove duplicates still resulted in similar completeness (BUSCO score of 95%) and an assembly size of over 11 Gb. Therefore, regardless of the assembly settings, the total length estimates confirm that the *Vampyroteuthis* genome constitutes the largest cephalopod genome sequenced to date (**Supplementary Figure 1)**. Since we could not currently obtain additional samples for this rare deep-sea species, chromosomal conformational data is not yet attainable. However, while our *Vampyroteuthis* draft genome assembly consists of numerous sub-chromosomal level contigs (about 5,600 contigs, N50 of 9.96 Mb, and the longest contig of 51.7 Mb), it still contains sufficient syntenic information^22^, with 162 contigs containing 15 or more orthologous genes and the largest contig containing 130 orthologous genes. Meanwhile, the size of the genome of *Argonauta hians* was ∼1.57 Gb, similar to that of its congener *A. argo*^33^, but only half of that of *Octopus* (∼3 Gb)^14^. The draft genome of *A. hians* was assembled into putative 28 chromosomes, comparable in contiguity and resolution to other octopod genomes, which typically have 30 chromosomes (**Supplementary Figure 2**).

Gene mixing on fused chromosomes is irreversible, and thus any ancestral or fusion-with-mixing (FWM) would serve as a strong synapomorphy for the clade in question^33,35,36^. To investigate how genome architectures relate to chromosomal evolution, we conducted comparative analyses across multiple cephalopod lineages using a modified synteny approach (see Methods). This approach enabled us to profile the presence of chromosomal irreversible FWM (**Figure 2**) and to confirm the highly rearranged nature of coleoid chromosomes relative to the nautilus (**Supplementary Figure 3**)^15^. Despite its gigantic size due to massive expansion of repetitive elements, we also found that the *Vampyroteuthis* genome structure predominantly adheres to basic coleoid karyotype rather than the more ancestral nautilus or the general molluscan chromosomal complements (**Supplementary Figure 3**)^22^. This pattern remained consistent even when alternative mapping and orthology search strategies were applied (**Supplementary Figure 4**), confirming the robustness of this result.

**Figure 2.**
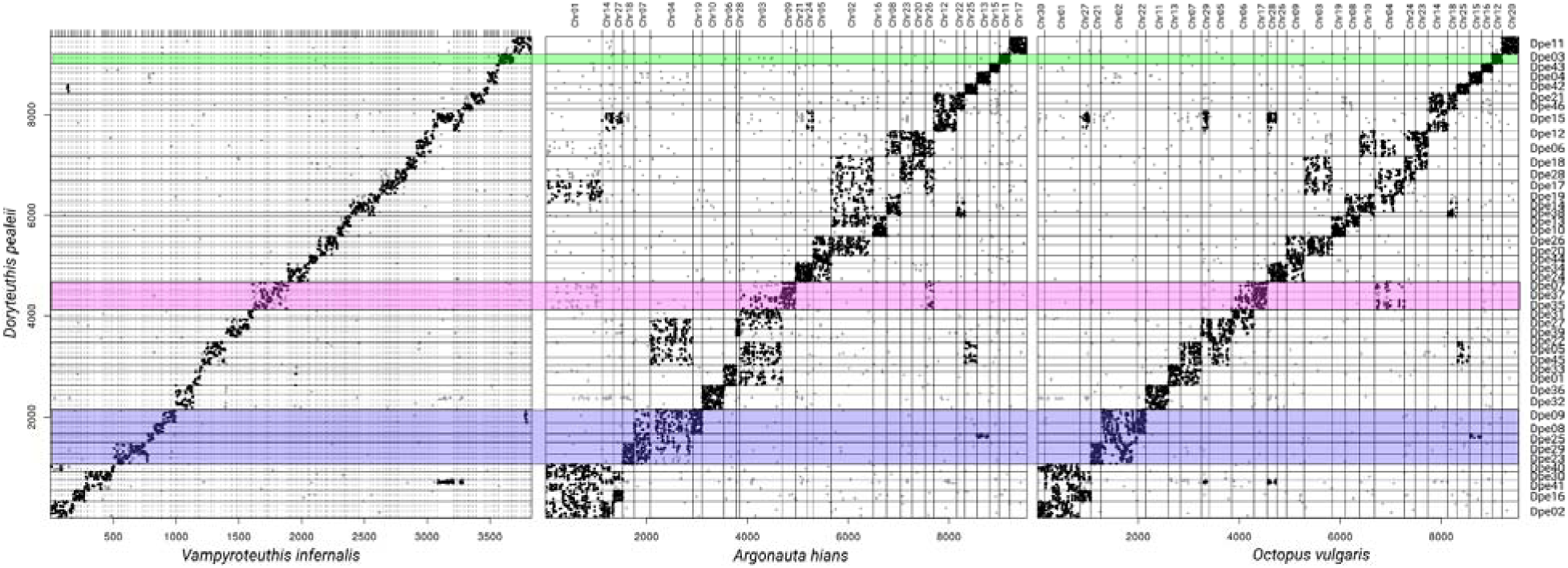
. *Vampyroteuthis* genome suggests ancestral coleoid karyotype was decapodiform-like, followed by karyotype reduction in Octopodiformes. Synteny dotplots showing Octopodiformes and *Argonauta-Octopus* shared FWM characters, relative to the *Doryteuthis pealeii* chromosomes. *Doryteuthis* chromosomes are ordered in the same way across the three species comparisons. The color codings indicate representative examples of the ancestral coleoid chromosome (green), ancestral Octopodiformes mixed chromosome (pink), and fused-and-mixed chromosomes in octopods (blue), respectively (corresponding to the phylogenetic nodes highlighted in Figure 1).

Surprisingly, we found that the *Vampyroteuthis* chromosomal structure is more closely aligned with that of Decapodiformes, including those of the longfin squid *Doryteuthis pealeii*, the bobtail squid *Euprymna scolopes*, the diamond squid *Thysanoteuthis rhombus*, and the cuttlefish *Sepia lycidas* (**Figure 2**, **Supplementary Figure 5**), despite its well-supported phylogenetic placement in Octopodiformes and thus its affinities with the octopods. Although our *Vampyroteuthis* draft genome assembly is not at the chromosomal level, its contigs often correspond to single *Doryteuthis* chromosomes, reflecting a high degree of chromosomal retention. Conversely, *Vampyroteuthis* contigs exhibited complex relationships with the chromosomes of the other octopodiform species (*A. hians*, the curled octopus *Eledone cirrhosa*, and the common octopus *Octopus vulgaris*), indicating synapomorphic fusions and rearrangements shared across octopods (**Figure 2**, **Supplementary Figure 5**). Multiple *Doryteuthis* chromosomes and their one-to-one homologous *Vampyroteuthis* sets of contigs correspond to single octopod chromosomes and show highly entropic mixing. For example, *O. vulgaris* chromosome 22 corresponds to *Doryteuthis* chromosomes 8 and 9, each with its own non-overlapping set of *Vampyroteuthis* contigs, with the degree of entropic mixing of these two states along the *Octopus* chromosome of 0.193, as measured by the normalized turbulence^37^. This implies that FWM of two ancestral decapodiform-like chromosomes (*Doryteuthis* chromosomes 8 and 9) occurred at the base of the octopod (*Eledone*, *Argonauta*, *Octopus*) lineage after its separation from *Vampyroteuthis*. For comparison, *O. vulgaris* chromosome 2 shows inter-chromosomal translocations with sharp syntenic boundaries and little mixing (<0.1 normalized turbulence for *Doryteuthis* chromosomes 9 and 29), suggesting a more recent origin of this translocation event. These chromosomal changes most likely accumulated gradually in the lineage leading to the octopods after their divergence from the *Vampyroteuthis* lineage (**Figure 2**, **Supplementary Figure 5**). Given the basal position of *Vampyroteuthis* among the Octopodiformes, the conservation of chromosomal structure across decapodiform lineages and *Vampyroteuthis*, along with the drastic chromosomal rearrangements shared by all octopods suggest the ancestral nature of the decapodiform karyotype.

To further explore the extent of chromosomal fusion and mixing in the lineage leading to octopods, we examined how *Doryteuthis* chromosomes were mapped onto the *Vampyroteuthis* genome. The results revealed chromosomal rearrangements with a high degree of mixing within the *Vampyroteuthis* contigs. For example, contig ptg000753l is mapped to *Doryteuthis* chromosomes 7, 37, and 35 with normalized turbulence measure of 0.285 for Dpe35 and Dpe37, 0.290 for Dpe07 and Dpe37, and 0.193 for Dpe07 and Dpe35^37^. The pattern is observed for 33 out of the 46 *Doryteuthis* chromosomes, where parts of the chromosomes were mapped onto the same set of *Vampyroteuthis* contigs (Fisher’s exact test p-value <0.05), suggesting irreversible fusion-with-mixing. As the same chromosomes were found to be fused and mixed in all other octopodiform species, this suggests an ancient synapomorphy supporting the affinity of *Vampyroteuthis* with octopods. On top of this synapomorphic signature, further substantial chromosomal fusion-with-mixing occurred in the lineage leading to octopods (*Eledone*, *Argonauta*, and *Octopus*) (**Figure 2**). Octopod homologs of only 3 *Doryteuthis* chromosomes (including the proposed sex chromosome^38^) remain unfused, suggesting that 43 out of 46 cases where chromosomes underwent fusions on the branch leading to octopods. Of these, at least 10 chromosomes were previously unfused in *Vampyroteuthis*, indicating additional fusions after the divergence of octopods from the vampyropod lineage. Comparison with *O. vulgaris* karyotype showed that the argonaut underwent further fusions between chromosomes 3 and 8, as well as 6 and 7, reducing the chromosome number from 30 to 28, suggesting a general trend of chromosomal number reduction in octopuses.

Our observations strongly suggest that the ancestral coleoid had a decapodiform-like karyotype, which later underwent additional fusions and karyotype reduction, first at the base of Octopodiformes, and then later in the octopods (**Figure 2**). The alternative scenario of chromosomal fissions is highly unlikely, as it would require independent, convergent fissions of hundreds of genes to produce identical chromosomal splits observed in all Decapodiformes and *Vampyroteuthis*^35^. We could also confirm this pattern by partitioning the *O. vulgaris* genome into fragments containing the same gene content as those contained in our *Vampyroteuthis* assembly. Using the same syntenic analysis approach, this artificially “rearranged and fragmented” *Octopus* genome was still recapitulating octopod fusions (**Supplementary Figure 6**), besides also showing a signal for the fusion of 42 out of 46 *Doryteuthis* chromosome homologs, compared to the 43 fused *Doryteuthis* chromosome homologs found in the complete *O. vulgaris* genome (Fisher’s exact test p-value <0.05). This suggests that the contiguity of the *Vampyroteuthis* assembly is unlikely to impact the inference of chromosomal fusions. These results suggest that *Vampyroteuthis* has a chromosomal structure partially shared with extant Decapodiformes, supporting the interpretation that it retains the more ancestral coleoid configuration, while octopods underwent additional FWMs.

We also observed a complex syntenic pattern in octopod chromosomes that cannot be explained by simple FWMs (**Figure 3**). Two possible scenarios might explain this complex syntenic pattern (**Figure 3b**). First, it may suggest that major inter-chromosomal translocations have occurred in the lineage leading to octopods (*Eledone*, *Argonauta*, and *Octopus*; **Figure 2b**). The second possibility is that a large post-ACCRE ancestral coleoid karyotype independently fused into different configurations in Decapodiformes and Octopodiformes (**Figure 3b**). If the second scenario was the case, we would expect to find a decapodiform species with an unfused chromosomal pair that corresponds to separate chromosomes in *Vampyroteuthis* or octopods yet fused in other Decapodiformes. However, since we have yet to find such decapodiform species, our data most strongly support the first scenario (**Supplementary Figure 7**). Although similar syntenic patterns can also arise after whole genome duplication^35^, we found no evidence of paralogous enrichment on any of the octopodiform chromosomes. Besides that, the observed chromosomal correspondence pattern does not indicate the presence of a consistent genome-wide doubling of ancestral chromosomes. Therefore, we believe it is unlikely for any whole-genome duplication event to have happened and be the cause of the syntenic patterns we observed. Additionally, comparisons within the octopods (**Supplementary Figure 8**) show that *Eledone*, despite its basal position on the tree, showed additional modifications to its karyotype, including translocations to the otherwise highly conserved ancestral coleoid chromosomes Dpe03-Ovu12. On the other hand, genomes of *A. hians* and *O. vulgaris* were strikingly similar and shared more chromosomal characteristics with *Vampyroteuthis*, suggesting a more ancestral octopod karyotype (**Supplementary Figure 8**).

**Figure 3.**
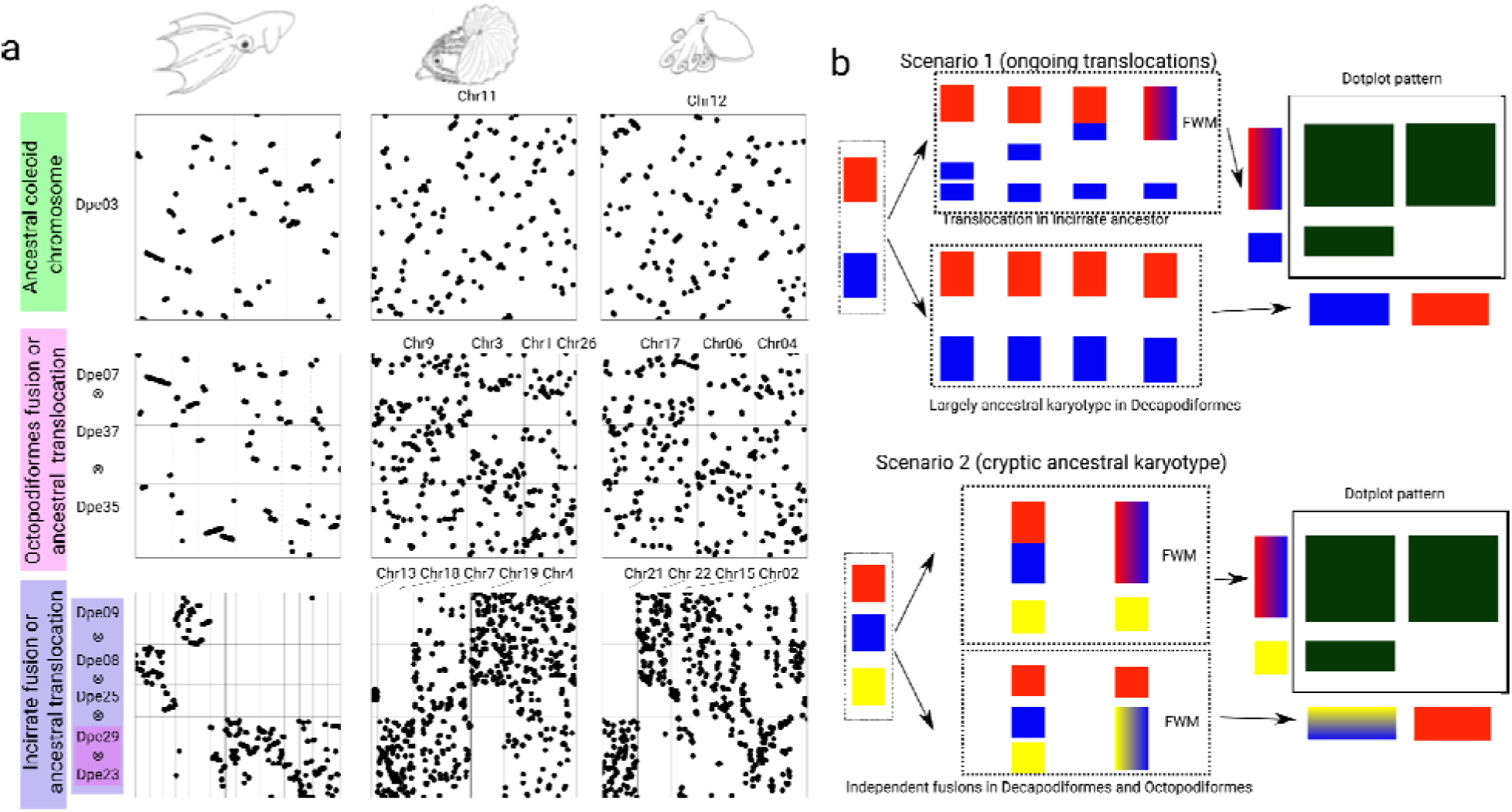
Chromosomal evolutionary pattern suggests additional ancestral translocations in octopods. (a) Dotplot representation for the three cases highlighted in Figure 1, with the Y axis indicating the chromosomes of *Doryteuthis*. Green indicates conserved coleoid chromosomes, as suggested by the conservation of genes in the chromosomes of all compared coleoids, despite translocation of the genes’ location within the chromosomes (intrachromosomal rearrangements; “mixing in a chromosome”). Meanwhile, ancestral Octopodiformes (pink) indicate fusions of different *Doryteuthis* chromosomes, which happened in ancestral Octopodiformes. The third example (blue) shows the formation of complex patterns of fusion-with-mixing, where orthologous genes undergoing intrachromosomal translocations in specific regions of the chromosomes of *Doryteuthis* are present in multiple chromosomes in compared species. (b) Two main scenarios for the observed complex chromosomal evolutionary patterns. The first one involves a fusion-with-mixing of intact (in red) and partial (in blue) chromosomes following inter-chromosomal translocations (blue) that occurred in Octopodiformes after divergence from the decapodiform karyotypes. A second possible scenario suggests that following the split of Octopodiformes and Decapodiformes, each lineage experienced distinct fusion-with-mixing events involving multiple ancestral chromosomes. In Octopodiformes, two specific chromosomes fused and mixed (red and blue), while in Decapodiformes, a different pair of chromosomes (blue and yellow) underwent a similar fusion-with-mixing process.

Why has the vampire squid maintained its more ancestral chromosomal state despite a dramatic increase in genome size? While several studies have implied the effects of transposable elements (TEs) accumulation on enhancing genome rearrangement rate, some of the largest genomes sequenced to date (e.g., lungfish^39^) show surprisingly well-conserved karyotypes. This suggests that the *Vampyroteuthis* genome provides another example where TE-driven genome expansion was decoupled from an increase in genome rearrangements. Future studies of both epigenetic states and genome topology in this species should provide fruitful insight into the role of TE accumulation in the maintenance of ancestral genomic configuration in animal genomes.

To further investigate the role of chromosomal fusion events onto putative regulatory landscapes as reflected by conserved non-coding element (CNE) preservation, we have conducted whole genome alignments between *Vampyroteuthis* and other coleoid cephalopods (Methods). We found that the conservation of non-coding regions was higher between *Vampyroteuthis* and Decapodiformes (over 7.1 Mb aligned sequences) compared to *Vampyroteuthis* and *Octopus* (2.5 Mb total alignment, **Figure 4a**). The decrease in the overall alignment length is particularly visible in *Argonauta* and *Octopus*. This pattern, while correlated with the genome size, is also corroborated by the overall repeat composition of the *Vampyroteuthis* genome, with long interspersed nuclear elements (LINEs) comprising a major part of the genome (14.12%) and short interspersed nuclear elements (SINEs) less than 1% (**Supplemental table 4**). This is similar to other Decapodiformes genomes and in contrast to the SINE-dominated octopod genomes^19^, indicating *Vampyroteuthis*’s higher propensity to retain ancestral non-coding features.

**Figure 4.**
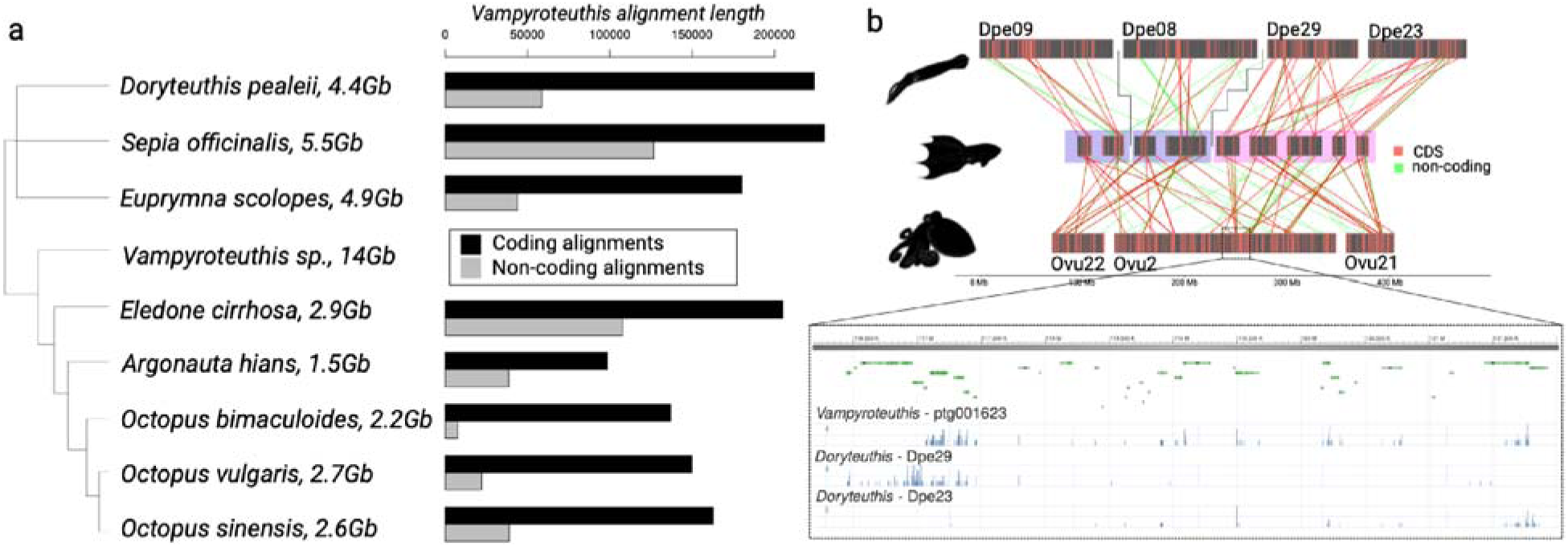
Conserved non-coding element complement in coleoid genomes. (a) *Vampyroteuthis*-centered whole genome alignments show the total content (in alignment numbers) of coding and non-coding alignments. (b) Ribbon diagram showing location of homologous alignments of coding (red, CDS) and non-coding (green) regions between *Doryteuthis* chromosomes (top) to *Vampyroteuthis* (middle) and *Octopus vulgaris* (bottom) for the complex FWM shown in Figures 2 and 3 (pink and blue colors). Zoom-in on a middle portion of *O. vulgaris* chromosome 2 (Ovu2) is highlighted below (NCBI genome browser) with gene track of annotation and conservation of both coding and non-coding elements derived from one homologous *Vampyroteuthis* contig and its pre-octopodiform unmixed state on two chromosomes Dpe29 and Dpe23 (pink color). Two *Vampyroteuthis* contigs homologous to Dpe09 and Dpe08, on the other hand, remain unmixed and their homologs undergo FWM only in octopods (lila color).

Using chromosomal homologies inferred from gene-level comparisons (**Figures 2** and **3**), we traced their CNE complement evolution on the derived octopod chromosomes. *Vampyroteuthis* homologs of unfused *Doryteuthis* chromosomes (Dpe09 and Dpe08) remain unmixed as separate units, whereas *Vampyroteuthis* contigs that correspond to the predicted octopodiform FWM events show substantial mixing of both coding and non-coding elements (Dpe29 and Dpe23, **Figure 4b**). All these contigs map to a set of *O. vulgaris* chromosomes, resulting in a high degree of mixing of gene loci and their putative regulatory sites. From this observation, we can infer that a large portion of the regulatory landscapes on modern *O. vulgaris* chromosomes has a complex evolutionary origin, shaped by FWM events combining segments derived from distinct chromosomal units in the ancestral octopodiform lineage.

## Discussion

Our findings show that the decapodiform-like karyotype represents the ancestral state of coleoids and support the basal position of *Vampyroteuthis* within Octopodiformes. This conclusion aligns well with paleontological data and insights regarding cephalopod bauplan evolution. Stem coleoid species from the Paleozoic, such as *Gordoniconus* and *Bactrites*, exhibit internal shells, streamlined morphologies, and ten arms, long before the appearance of the oldest vampyropods in the Jurassic, such as *Teudopsis* and *Vampyronassa*. Interestingly, *Vampyroteuthis* also possesses two long feeding filaments, thought to be vestigial arms, indicating a possible morphological affinity to the 10-armed decapodiform. The retention of parts of the decapodiform-like chromosomal architecture in *Vampyroteuthis*, along with the extensive chromosomal rearrangements observed in octopods, provides genomic evidence supporting the transition from decapodiform-like ancestors to Octopodiformes. Our analyses also indicate a surge of inter-chromosomal translocations at the base of the octopods, potentially facilitated by demographic shifts or genomic instability. These ACCRE-like inter-chromosomal translocations, possibly stabilized by the notably large genome sizes in coleoids, may have driven morphological innovations and regulatory complexity. For instance, the specialization of arms into feeding tentacles in squids and the reduction of arms in octopuses likely result from modifications in gene regulation and its chromosomal context, and not gene content. Similarly, the presence of shell matrix protein-coding genes in shell-less octopods suggests that regulatory changes, rather than structural ones^33,40^, facilitated shell degeneration. Fossil record and molecular data provide a temporal framework to time of these innovations (**Figure 1a**), including the accumulation of chromosomal rearrangements and their impacts on phenotypic evolution in the octopod lineage.

Taken together, our findings have important implications for understanding the directionality of chromosomal evolution in the animal kingdom, its long-term impact on the emergence of novel putative regulatory regions, and, eventually, morphological innovations. Further attempts to obtain a chromosomal-scale assembly of *Vampyroteuthis* will help to determine the exact karyotypic changes that occurred at the base of Octopodiformes.

## Materials and Methods

### Samples

The genomes of a single juvenile *Vampyroteuthis* of unknown sex, caught in Suruga Bay by T/V Hokuto of Tokai University, and a single *Argonauta hians* adult female caught as bycatch in fixed nets set along the coasts of Oki Island, Shimane Prefecture, Japan (36°17′20.6″ ″N 133°12′46.4″″E), were sequenced. The eggcase of sequenced *A. hians* specimen was deposited in the University Museum, the University of Tokyo (UMUT RM34224).

### Sample preparations

Whole-genome shotgun sequencing was performed using PacBio and Illumina sequencing platforms. Genomic DNA was extracted from the muscles (whole body) of *Vampyroteuthis* and the ovary of *A. hians* using a Genomic-tip Kit (QIAGEN, Hilden, Germany) and then sheared into fragments (size: 15 kb to 20 kb) with a g-tube device (Covaris Inc., MA, USA). PacBio HiFi libraries for *Vampyroteuthis* and *A. hians* were prepared using a SMRTbell Prep Kit 3.0 (Pacific Bioscience, CA, USA) according to the manufacturer’s instructions, and were size-selected using the SageELF system (Saga Science, MA, USA). Seventeen SMRT cells for *Vampyroteuthis* libraries and two SMRT cells for *A. hians* libraries were sequenced on the PacBio Sequel II/IIe systems with Binding Kit 3.2 and Sequencing Kit 2.0 (30 hours collection times). The consensus (HiFi) reads were generated from raw full-pass subreads using the DeepConsensus v1.1.0 program^41^ with default parameter settings. For Illumina sequencing, genomic DNA was fragmented to an average size of 500 bp using the Focused-ultrasonicator M220 (Covaris Inc., MA. USA). Paired-end libraries were constructed with a TruSeq DNA PCR-Free Library Prep kit (Illumina, CA, USA) and size-selected on an agarose gel with a Zymoclean Large Fragment DNA Recovery Kit (Zymo Research, CA. USA). The final libraries were sequenced on a NovaSeq 6000 system (Illumina, San Diego, CA, USA) with 2 × 150 bp read length.

Total RNA was extracted from three *Vampyroteuthis* tissues (brain, buccal mass, and eyes) and four *A. hians* tissues (brain, eyes, heart, and embryo) using E.Z.N.A. Mollusc RNA kit (Omega Bio-Tek, GA. USA) and a Nucleospin RNA clean-up XS (TaKaRa, Japan). We obtained the tissues from the same individuals to the genome DNA. The RNA-seq libraries were constructed using an Illumina Stranded mRNA Prep, Ligation (Illumina, San Diego, CA, USA) following the manufacturer’s protocol. Sequencing was conducted on the NovaSeq 6000 system with 2 x 100 bp read length. Library concentration and qualities were assessed using Qubit 4 Fluorometer (Thermo Fisher Scientific, MA, USA), the 2100 Bioanalyzer system (Agilent Technologies, CA, USA), and 7900HT Fast Real-Time PCR System (Thermo Fisher Scientific, MA, USA).

### Genome sequencing and informatics

In total, 571,504,779,822 bp HiFi reads were obtained by PacBio Sequel II/IIe for *Vampyroteuthis*. The total sequencing depth was approximately 40X, and the average read length was 16.3 kb. 1,303,449,174,900 bp of clean data were acquired through illumina NovaSeq6000 (PE500) short reads. We analyzed the whole genome by Hifiasm^42,43^, followed by purge_dups^44^. The details of the assembly, repeat modelling, and annotation are presented in the supplementary material.

Genome assembly of *A. hians* was conducted by Hifiasm using standard parameters. Hi-C libraries for scaffolding were constructed using Dovetail Omni-C Kits (Cantata Bio, CA, USA), and were sequenced on an Illumina MiSeq system. The Hi-C reads were mapped to the contigs and filtered using Juicer v1.6^45^. Scaffolding was performed with 3D-DNA v180419^46^. The resulting scaffolds were manually inspected and refined using Juicebox v1.11.08^47^.

### Orthology and syntenic analysis

For consistency in orthology handling, we mapped *Doryteuthis pealeii* and *Octopus vulgaris* peptides to all genomes using miniprot 0.12^48^ using-gff option as an output. In total, 30,432 *D. pealeii* and 25,682 *O. vulgaris* genes were mapped. Custom scripts were used to parse the mapping results and convert them into pairwise synteny files (.psynt) for each species pair (available on the repository link below). Each .psynt file contains information on mutual best orthologs as well as their genomic locations in two species. Only contigs with 15 or more orthologous genes were used. Dotplots were made using a custom R (version 4.2.2) script psyntPlot.R using published procedures^49^. Significance of chromosomal homologies was tested using Fisher’s exact test, and associations of adjusted p-value (Bonferroni correction) of 0.05 are shown as black colored dots. Clustering of contigs or chromosomes on dotplots was done either by predefined order of chromosomes (Figure 2) to enable multi-species comparisons, or via Euclidean distance measure on the number of shared orthologs and ward.D2 clustering in R. Statistical analysis of mixing was done with TraMineR 2.2-11 package in _R_48_._

Whole genome alignments were done on hard-masked genomes (RepeatModeler 2.0.6 and RepeatMasker 4.1.8) with blastn 2.16.0+ using the following parameters: -task megablast-perc_identity 0-template_length 16 -penalty -2 - word_size 11-evalue 1-template_type coding_and_optimal^50^. Transdecoder 5.7.1 was used to predict open reading frames on conserved aligned sequences (TransDecoder.LongOrfs-m10 parameter was used) and classify alignments into coding vs non-coding regions, in addition to available gene annotation overlap. Only alignments of 50 or more base pairs were reported. All dotplots for every figure and their supporting data can be generated by running the prepInput.sh and plot_figures.R scripts on the repository.

## Acknowledgments

We thank Noriyoshi Sato (Tokai University) for providing *Vampyroteuthis* samples. We also thank all members of the Simakov lab (University of Vienna), in particular Thea Rogers and Darrin Schultz, for invaluable discussions and comments on the manuscript. D.H.E.S. and K.H. would like to thank Nanami Tochino (Setiamarga Lab at NIT Wakayama), for her assistance on the phylogeny/timetree inference.

## Author contributions

This work is a result of an equal collaboration among three labs led by three Principal Investigators (the corresponding authors: M.A.Y., D.H.E.S. and O.S.), all giving their shares equally toward the completion of this project. M.A.Y., D.H.E.S. and O.S. designed and led the study; M.A.Y., A. T., H. T., and H. N. contributed to the genome sequencing and data deposit; E.T., K.K., T. K., K. H., H. M., M. T., and H. N. conducted the bioinformatical work; M.A.Y., D.H.E.S. and O.S. produced the first draft of the manuscript. All authors contributed to the final manuscript.

## Supplemental information

Figures S1–S8, Table S1-S4

