## Supplement for "Giant genome of the vampire squid reveals the derived state of modern octopod karyotypes"

### **Supplemental note for the genome assemblies**

The PacBio libraries were constructed using the genomic DNA from the *Vampyroteuthis* sample. In total, 571,504,779,822 bp of HiFi reads were obtained by PacBio Sequel II/IIe. The total sequencing depth was approximately 40X, and the average read length was 16.3 kb. Short reads by illumina NovaSeq6000 (PE500) were also obtained, and 1,303,449,174,900 bp of clean data were acquired. Two RNA-seq libraries were obtained from the same individuals of *Vampyroteuthis*.

We assembled the whole genome by Hifiasm^1,2^ with an option --hg-size 14g since preliminary k-mer analysis showed a genome size of 14 Gb (**Supplemental Table 1**). We note that the results of the vampire squid stably reproduced 14G assembly size regardless of options. The v1.0 assembly we showed has a genome size of 14.7 Gbp and an N50 length of 7.6 Mbp. The genome size was 14.5 Gbp and N50 length 7.8 Mbp as a result of option 12G specification (hifiasm --hg-size 12g). Even using the latest version of hifiasm, the genome size was 14.4 Gbp and N50 length 8.5 Mbp with the 12G designation, and the N50 was extended a little further (hifiasm-0.24.0-r702 --hg-size 12g, 2024Dec_ver). Thus, regardless of the genome size setting option, the final assembly size remains the same across the currently available methods. BUSCO analysis showed that its completeness was as high as 97.2. This score was comparable to other chromosome-level Molluscan genomes (**Supplemental Table 2**).

K-mer based genome size estimation was applied by GenomeScope2.0^3^ (-k 35 -p 2). Jellyfish^4^ run was performed with an option: high 10000000. The estimated genome size was 11.5 Gb, and heterozygosity was estimated to be 1.8-2% (**Supplemental figure 1**). Another k-mer condition (k=27) gave almost the same result (11.8 Gb). The Hifiasm standard output size is significantly larger than the genome size estimated by k-mer analysis of short-read data. The discrepancy in assembly size is probably the result of duplication due to haplotype diversity. BUSCO analysis also detected a high number of duplicate genes (**Supplemental Table 3**). Based on these findings, contig duplication was suspected. We applied purge_dups to reduce the duplicated content to a haplotype consensus sequence. The purged assembly data (v2.0) set has a size of 11.7 Gb and 5,616 contigs with an N50 contig size of 9.96 Mbp. GenomeScope 2.0 estimated an expected genome size of 11.5-11.8 Gb, which is close to the assembled size after applying purge_dups. The purge_dups collapses the haplotype-separated assembly and reduces the duplicated content to a haplotype consensus sequence, therefore, contigs purged in this process are probably duplicated contigs due to haplotype diversity. The purged contigs are not enriched with a high mask rate and appear to be a population of contigs derived from minor haplotigs with the same distribution as the whole (**Supplemental figure 1**). This is also in good agreement with the improved duplicate comparing BUSCO scores before and after purge_dups (7.1% to 3.5%). For *Argonauta hians*, GenomeScope 2.0 estimated an expected genome size of 1.56 Gbp, which is consistent with the assembled genome size (1.57 Gbp; version 1.0).

Repetitive elements of the *Vampyroteuthis* genome were estimated with Repeatmodeler^5^. However, LTR structural analysis (LtrHarvest::gt command) was not finished for over a month, probably due to the large-sized genome. So, we applied an alternative method; the LTRs that were not detected by repeatmodeler (-LTRStruct off), and additionally estimated using ltrDetector^6^. The query was compared to the classified sequences in "rpmd_db2-families.fa" using RepeatMasker version 4.1.2-p1 (default mode) with rmblastn version 2.11.0+. This method is practical and finished within a day. 62.47% of the genome was estimated as repetitive regions and masked for the following analysis (**Supplemental Table 4**). For *A. hians*, a repeat mask was followed by an estimation of the gene model. The repeat libraries were generated using RepeatModeler v2.0.5^5^ with RECON v1.08^7^, RepeatScout v1.0.6^8^, and RMBlast v2.9.0 (http://www.repeatmasker.org/RMBlast.html). Long terminal repeat sequences were also identified with “-LTRStruct” option, using LtrHarvest v1.5.9^9^ and Ltr_retriever v2.6^10^. Repeat annotation was performed with RepeatMasker v4.1.5^11^ using the custom libraries.

Gene prediction models were generated using GINGER 1.0.1^12^. In brief, this pipeline combines *ab initio*, RNA-seq-based, and homology-based prediction results for related species. For homology-based prediction, the amino acid sequences of *Octopus vulgaris*^13^, *O. bimaculoides*^14^, *Architeuthis dux*^15^, *Crassostrea gigas*^16^, and *Mizuhopecten yessoensis*^17^ were utilized and mapped to the genome scaffolds. The predicted genes were evaluated using BUSCO^18^, and resulted in 96.6% complete genes being marked, suggesting high accuracy of the annotation [C:96.6%[S:93.9%,D:2.7%],F:1.9%,M:1.5%,n:954]. The initial gene prediction using Ginger yielded a total of 82,025 gene models. This number is extremely high compared to other octopus species, likely due to the difficulty of accurately masking repeat sequences in large genomes. To address this, we applied a series of filtering steps to remove spurious gene predictions. Specifically, we excluded genes overlapping with LTRs identified by ltrDetector but not masked by RepeatMasker, and we removed genes lacking homology to related species, identifiable protein domains, or RNA-seq support. Although these filters significantly reduced the total gene count, the BUSCO completeness score remained unchanged. The final gene set for the Vampyroteuthis genome includes 62,225 protein-coding genes.

For *A. hians*, PacBio library was constructed using the genomic DNA from a single individual. In total, 94,777,318,231 bp of HiFi reads were generated by PacBio Sequel II/IIe. In addition, short-read sequencing was performed on the same sample using the Illumina NovaSeq 6000 platform (PE500), yielding 167,870,168,100 bp of clean data. of clean data. Four RNA-seq libraries were obtained from the same individuals of *A. hians*. De novo genome assembly was conducted using Hifiasm with default parameters, resulting in an initial assembly of 389 contigs with an N50 of 29.3 Mb and a maximum contig length of 130.8 Mb. This assembly was further scaffolded using Omni-C data.Hi-C library preparation, scaffolding, and gene model prediction were performed as described in the main text.

**Supplemental Table 1. Starts comparison genome assemblies shown in this paper**

| Genome version | **Vampyroteu-this sp v1.0** | **Vampyroteu-this sp v2.0** | **A. hians v1.0** | **A. hians v2.0** |
| --- | --- | --- | --- | --- |
| Difference from other versions | hifiasm standard output | duplicates purged using purge_dups | hifiasm standard output | Hi-C scaffolding |
| Total nucleotides | 14,690,897,198 | 11,710,348,364 | 1,636,068,110 | 1,517,914,136 |
| Number of contigs | 12,099 | 5,616 | 389 | 28 |
| N50 | 7,566,590 | 9,963,774 | 30,166,387 | 60,915,548 |
| L50 | 490 | 326 | 18 | 5 |
| Longest contig | 51,721,974 | 51,721,974 | 138,080,145 | 223,252,000 |
| Average length | 1,214,224.08 | 2,085,175.99 | 4,205,830.62 | 54,211,219.14 |

**Supplemental Table 2. Starts comparison cephalopod genomes**

|  | genome size | n_proteins | busco proteins mode  (metazoa_odb10, n:954) |
| --- | --- | --- | --- |
| *Vampyroteuthis infernalis* (v1.0) | 14.6 gb | 88,329 | C:97.2%[S:91.2%,D:6.0%],F:2.0%,M:0.8% |
| *Argonauta argo* | 1.3 gb | 20,293 | C:95.4%[S:93.1%,D:2.3%],F:2.6%,M:2.0% |
| *Architeuthis dux* | 2.7 gb | 51,225 | C:85.5%[S:84.8%,D:0.7%],F:7.5%,M:7.0% |
| *Crassostrea gigas* | 557 mb | 51,045 | C:95.4%[S:70.8%,D:24.6%],F:0.3%,M:4.3% |
| *Mizuhopecten yessoensis* | 971 mb | 41,567 | C:98.6%[S:75.2%,D:23.4%],F:0.4%,M:1.0% |
| *Octopus bimaculoides* | 2.3 gb | 29,037 | C:95.2%[S:69.8%,D:25.4%],F:2.3%,M:2.5% |
| *Octopus vulgaris* | 2.7 gb | 30,134 | C:91.2%[S:67.2%,D:24.0%],F:3.7%,M:5.1% |

**Supplemental Table 3. BUSCO statistics showed the eliminations of duplicated contigs by the purge_dups**

|  | **Original**  **assembly** | **After purge_dups applied** |
| --- | --- | --- |
| version | 1.0 | 2.0 |
| genome size | 14.6 Gb | 11.7 Gb |
| N_contigs | 12,095 | 5,616 |
| repeats | 60.8% | 60.3% |
| busco  (genome mode,  vs metazoa_odb10,  n:954) | C:95.8%  [S:88.7%,  D:7.1%],  F:3.1%,  M:1.1% | C:95.3%  [S:91.8%,  D:3.5%],  F:3.2%,  M:1.5% |

**Supplemental Table 4. Repeat content of the Vampyroteuthis genome (v.2.0) based on Repeat Masker output**

|  |  | number of elements | Length occupied (bp) | Percentage |
| --- | --- | --- | --- | --- |
| Retroelements |  | 2,914,604 | 1,979,850,998 | 16.91 % |
| SINEs: |  | 83,745 | 9217235 | 0.08% |
| Penelope |  | 563,170 | 300,260,284 | 2.56% |
| LINEs: |  | 2,468,426 | 1,653,598,627 | 14.12% |
|  | CRE/SLACS | 3,647 | 2,242,901 | 0.02% |
|  | L2/CR1/Rex | 851,669 | 530,457,586 | 4.53% |
|  | R1/LOA/Jockey | 34,896 | 14,318,566 | 0.12% |
|  | R2/R4/NeSL | 278,491 | 183,054,340 | 1.56% |
|  | RTE/Bov-B | 150,980 | 71,897,444 | 0.61% |
|  | L1/CIN4 | 12,271 | 6,622,898 | 0.06% |
| LTR elements: |  | 362,433 | 317,035,136 | 2.71% |
|  | BEL/Pao | 4,627 | 5,326,317 | 0.05% |
|  | Ty1/Copia | 2,264 | 5,727,452 | 0.05% |
|  | Gypsy/DIRS1 | 345,898 | 301,359,837 | 2.57% |
|  | Retroviral | 6,119 | 1,190,412 | 0.01% |
| DNA transposons |  | 1,931,104 | 724,641,702 | 6.19% |
|  | hobo-Activator | 746,706 | 280,232,486 | 2.39% |
|  | Tc1-IS630-Pogo | 617,804 | 247,553,787 | 2.11% |
|  | PiggyBac | 70,298 | 21,725,655 | 0.19% |
|  | Tourist/Harbinger | 11,518 | 1,969,450 | 0.02% |
|  | Other (Mirage, P-element, Transib) | 5,972 | 3,246,023 | 0.03% |
| Rolling-circles |  | 131,303 | 38,104,061 | 0.33 % |
| Unclassified: |  | 25,981,963 | 4,611,039,671 | 39.38 % |
| Total interspersed repeats: |  |  | 7,315,532,371 | 62.47 % |
| Small RNA: |  | 123,779 | 36,746,815 | 0.31 % |
| Satellites: |  | 10 | 8,614 | 0.00% |
| Simple repeats: |  | 5,286,058 | 459,293,181 | 3.92 % |
| Low complexity: |  | 250,904 | 20,431,074 | 0.17 % |

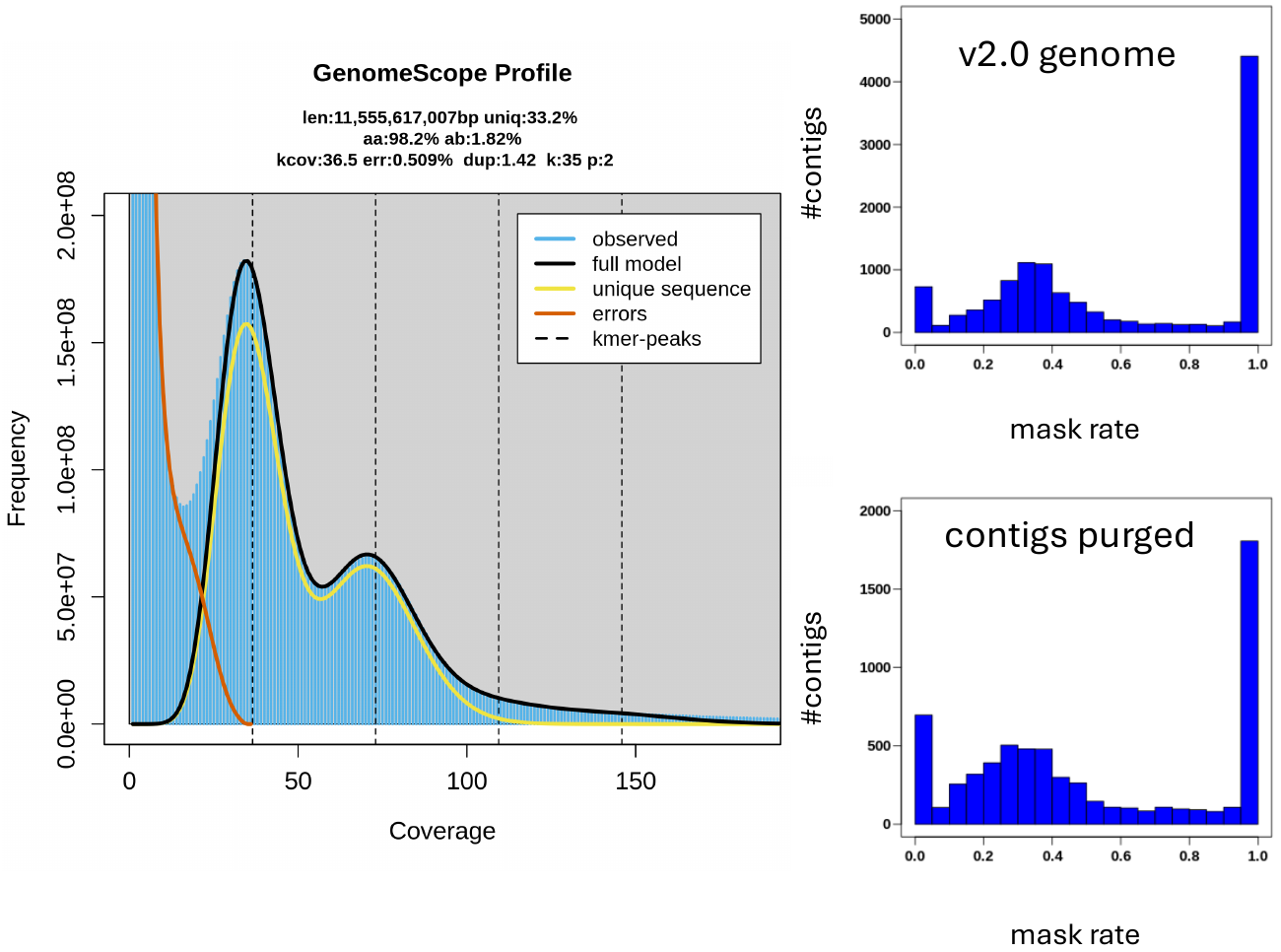
Supplementary Figure 1. **Genome size estimation via GenomeScope2.0.** Estimated results at k=35 are shown; estimates at k=27 were almost identical. Heterozygosity rate was 1.82%. Verification of possible existence of overlapping contig due to high heterozygosity. The assembly size of the ver 2.0 genome is close to the estimated genome size. The purged contig is not enriched with a high mask rate. There is a possibility that the current assembly contains duplicated contigs.

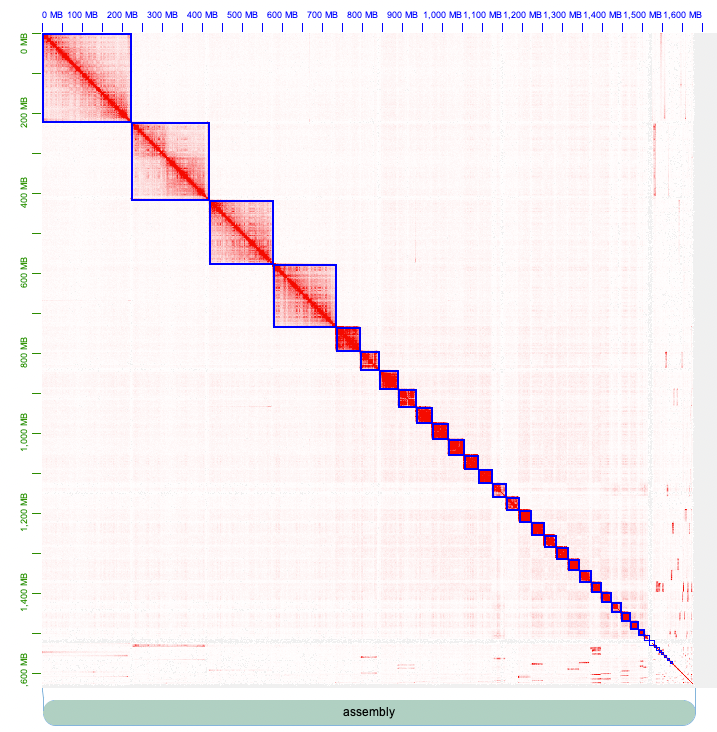

Supplementary Figure 2. **Argonauta hians HiC map**.

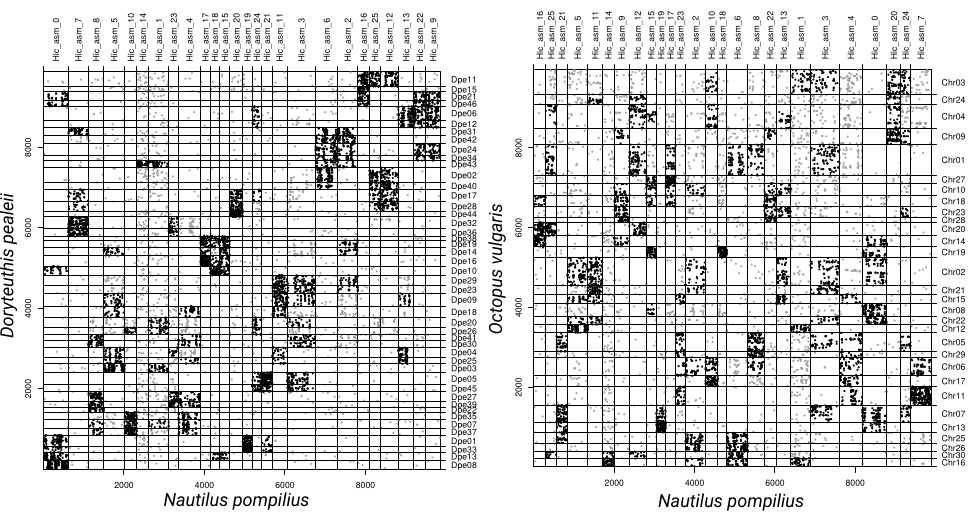

Supplementary Figure 3. **Translocation-rich history in coleoid genomes.** *Octopus vulgaris* and *Doryteuthis pealeii* genomes show many inter-chromosomal translocations and fusion-with-mixings compared to the Nautilus genome, which represents the ancestral molluscan karyotype. Significant associations (Fisher’s exact test p-value <0.05) are labelled in bold color.

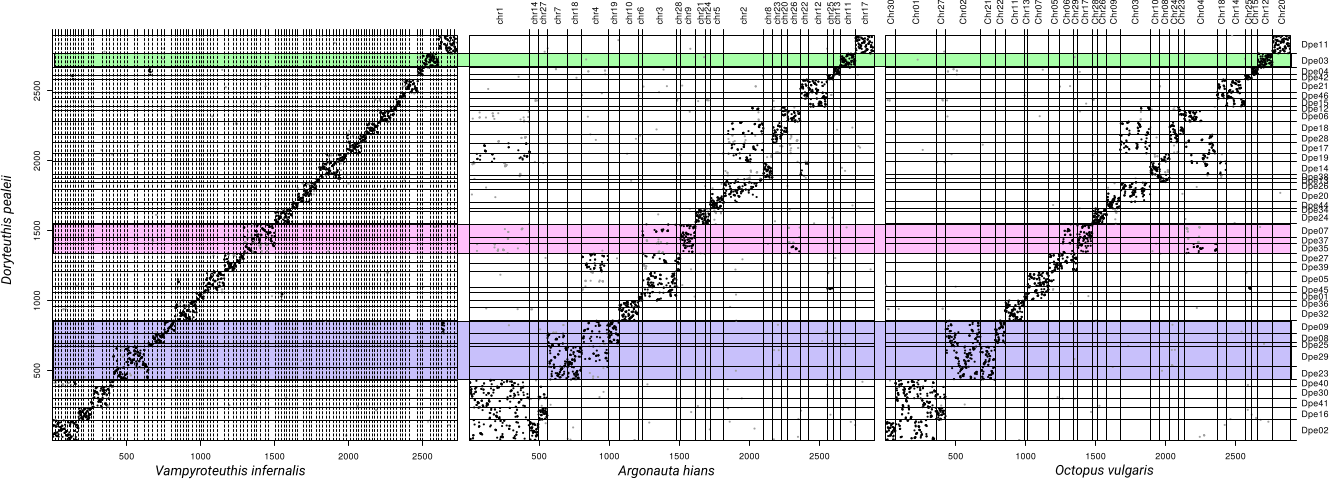

Supplementary Figure 4. **Syntenic pattern using a different orthology set (O. vulgaris gene annotations as reference)**. Same arrangements as in Figure 2.

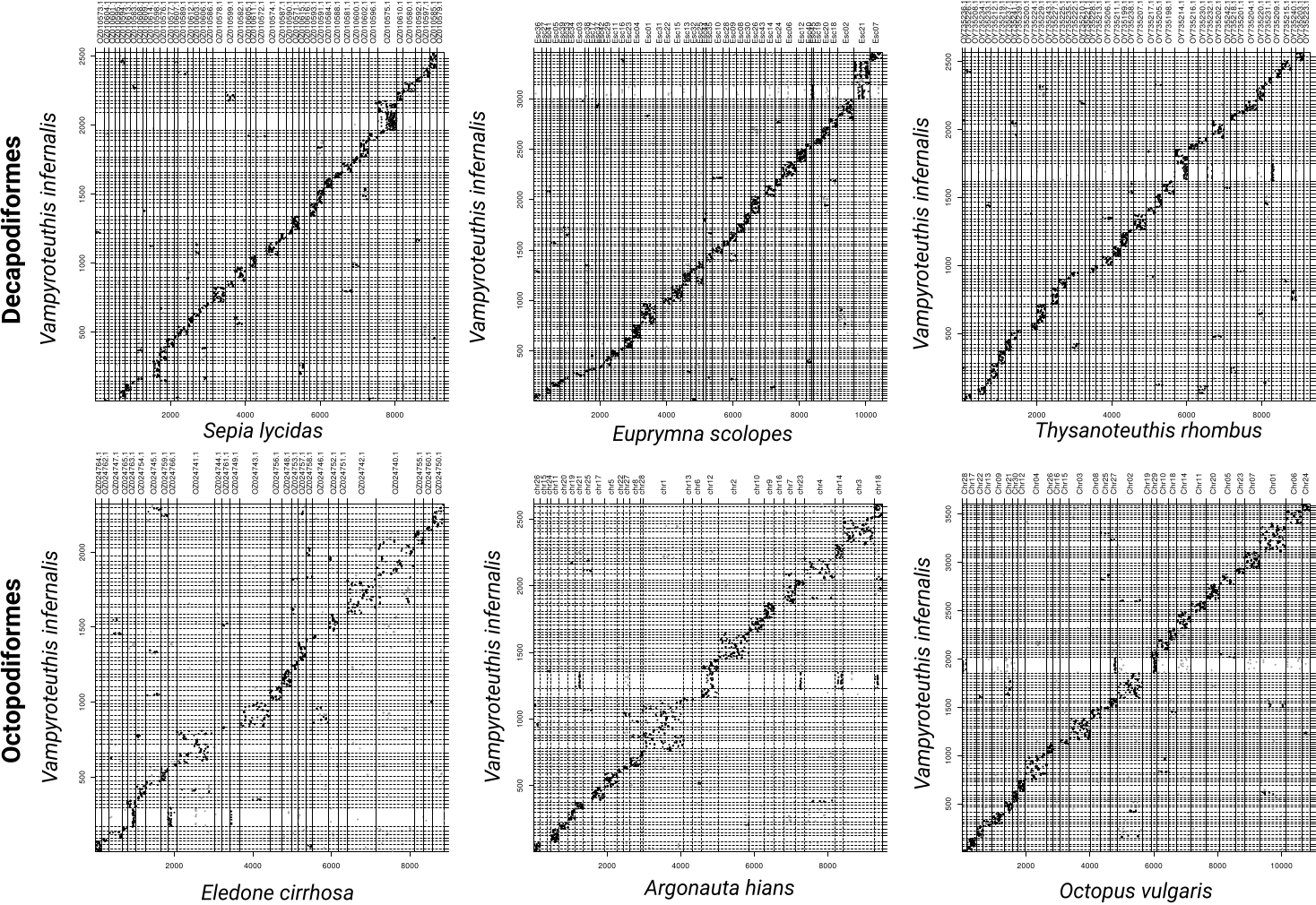
Supplementary Figure 5. ***Vampyroteuthis* shows an intermediate karyotype between Octopodiformes and Decapodiformes.** Significant associations (Fisher’s exact test p-value <0.05) are labeled in bold color.

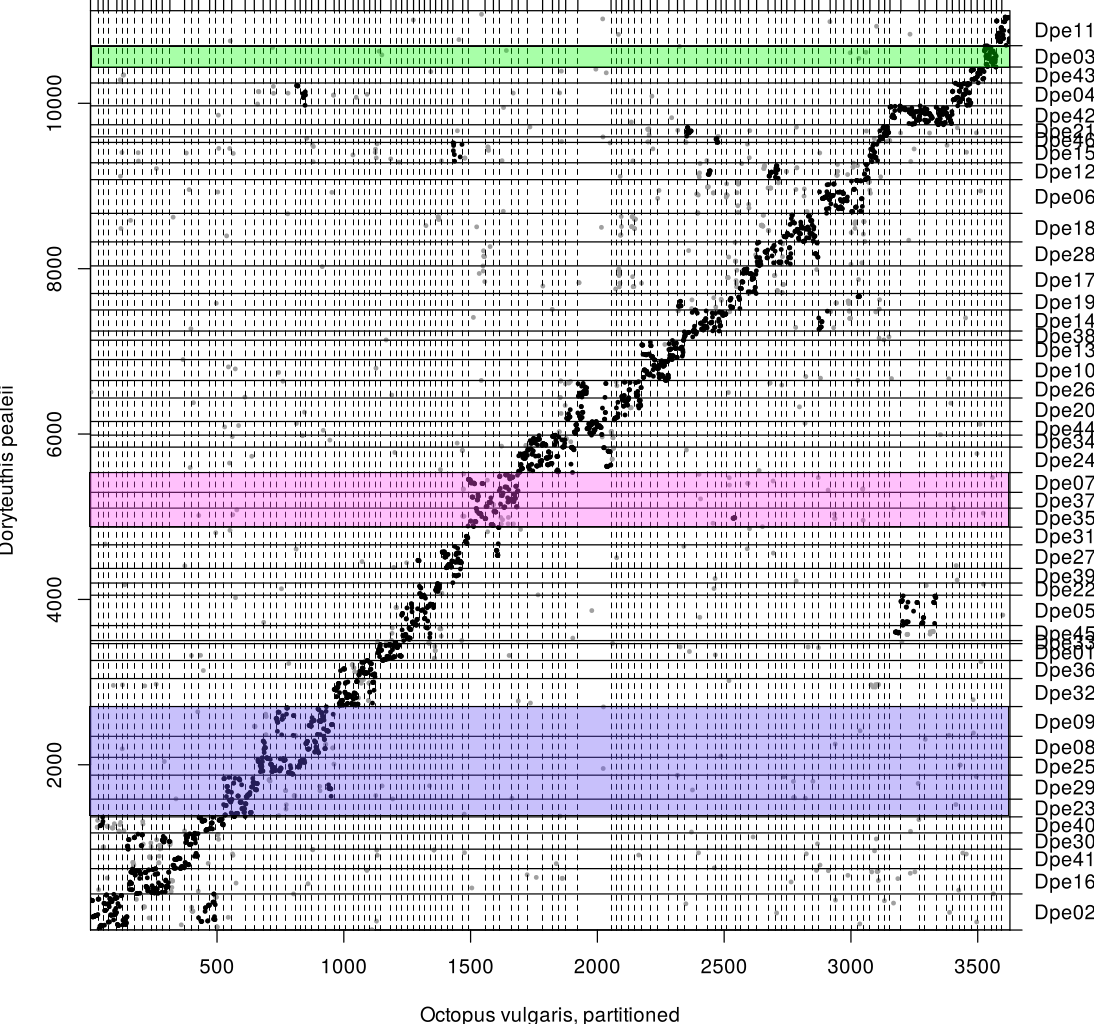
Supplementary Figure 6. ***Octopus vulgaris* genome, broken up into pieces of similar size as *Vampyroteuthis* contigs, recapitulates FWMs seen with chromosomal-level assembly.** Significant associations (Fisher’s exact test p-value <0.05) are labeled in bold color.

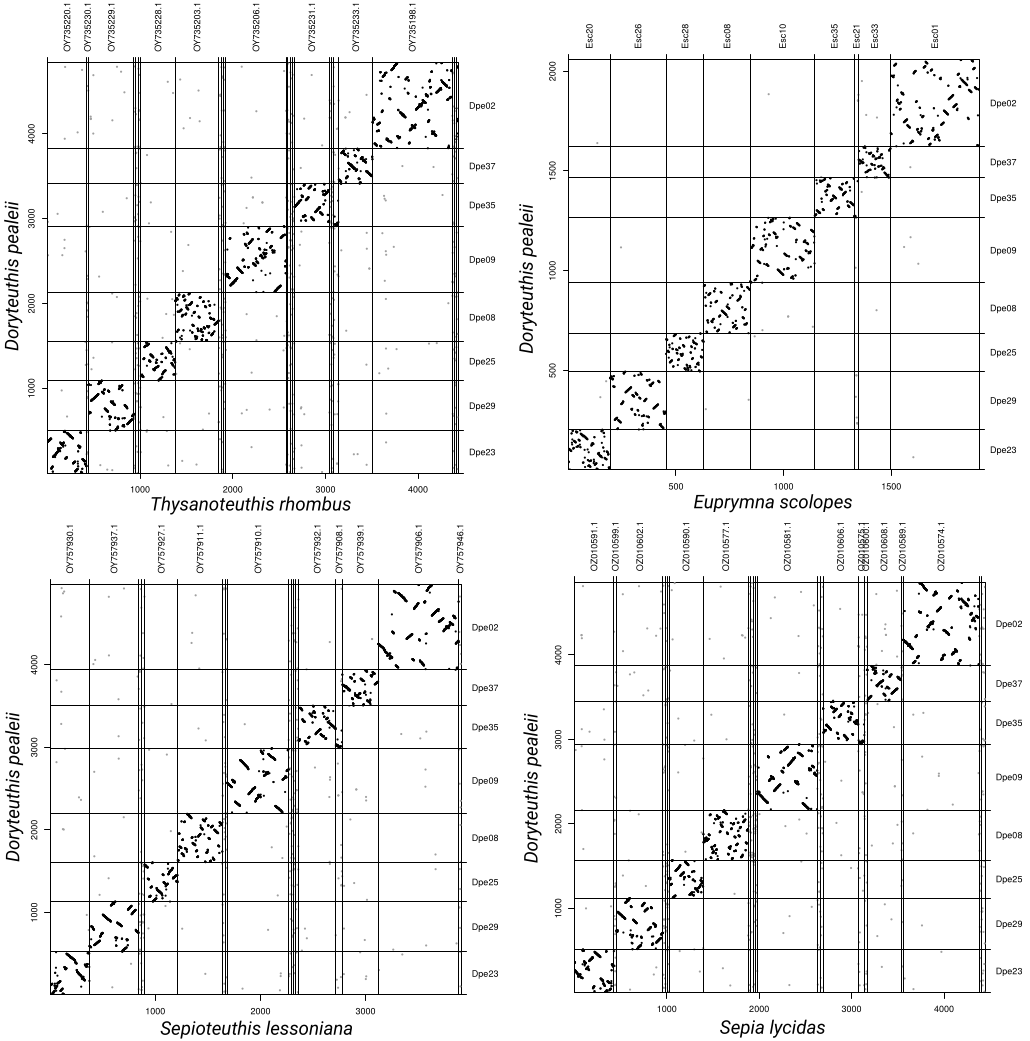
Supplementary Figure 7. **Decapodiformes chromosomes are conserved across their major lineages.** Dotplots for the selected (Figure 2) *Doryteuthis* chromosomes are shown, highlighting the absence of any “cryptic” chromosomal homologies (all chromosomes show 1-1 homology) in comparison with pelagic squids, as well as a bobtail squid and a cuttlefish. Significant associations (Fisher’s exact test p-value <0.05) are labeled in bold color.

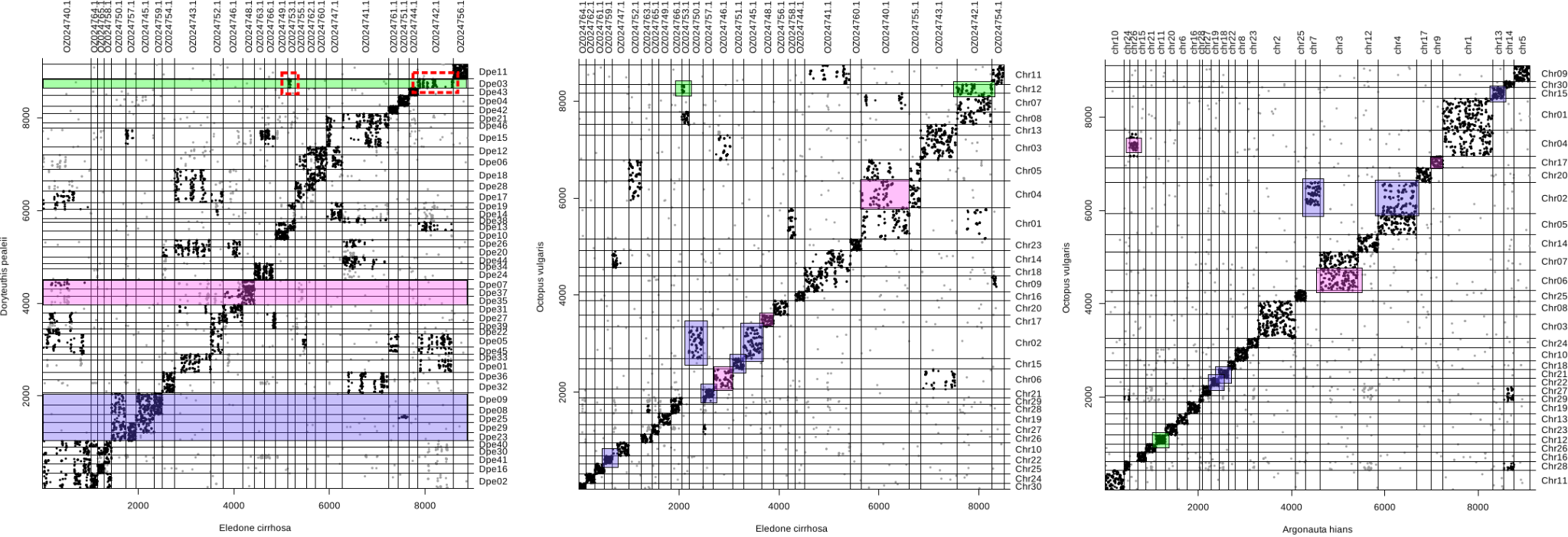

Supplementary Figure 8. **Eledone shows a more derived karyotype**. Dotplots within octopod genomes highlight high conservation between *A. hians* and *O. vulgaris*. Eledone, on the other hand, has additional translocations with one ancestral coleoid unit (Dpe03-Ovu12) split across multiple chromosomes. Color scheme as on Figures 2 and 3.
